# Sex-Specific Deflection of Age-Related DNA Methylation and Gene Expression in Mouse Heart by Perinatal Toxicant Exposures

**DOI:** 10.1101/2024.04.25.591125

**Authors:** Kai Wang, Maureen A. Sartor, Justin A. Colacino, Dana C. Dolinoy, Laurie K. Svoboda

## Abstract

**Background:** Global and site-specific changes in DNA methylation and gene expression are associated with cardiovascular aging and disease, but how toxicant exposures during early development influence the normal trajectory of these age-related molecular changes, and whether there are sex differences, has not yet been investigated.

**Objectives:** We used an established mouse model of developmental exposures to investigate the effects of perinatal exposure to either lead (Pb) or diethylhexyl phthalate (DEHP), two ubiquitous environmental contaminants strongly associated with CVD, on age-related cardiac DNA methylation and gene expression.

**Methods:** Dams were randomly assigned to receive human physiologically relevant levels of Pb (32 ppm in water), DEHP (25 mg/kg chow), or control water and chow. Exposures started two weeks prior to mating and continued until weaning at postnatal day 21 (3 weeks of age). Approximately one male and one female offspring per litter were followed to 3 weeks, 5 months, or 10 months of age, at which time whole hearts were collected (n ≥ 5 per sex per exposure). Enhanced reduced representation bisulfite sequencing (ERRBS) was used to assess the cardiac DNA methylome at 3 weeks and 10 months, and RNA-seq was conducted at all 3 time points. MethylSig and edgeR were used to identify age-related differentially methylated regions (DMRs) and differentially expressed genes (DEGs), respectively, within each sex and exposure group. Cell type deconvolution of bulk RNA-seq data was conducted using the MuSiC algorithm and publicly available single cell RNA-seq data.

**Results:** Thousands of DMRs and hundreds of DEGs were identified in control, DEHP, and Pb-exposed hearts across time between 3 weeks and 10 months of age. A closer look at the genes and pathways showing differential DNA methylation revealed that the majority were unique to each sex and exposure group. Overall, pathways governing development and differentiation were most frequently altered with age in all conditions. A small number of genes in each group showed significant changes in DNA methylation and gene expression with age, including several that were altered by both toxicants but were unchanged in control. We also observed subtle, but significant changes in the proportion of several cell types due to age, sex, and developmental exposure.

**Discussion:** Together these data show that perinatal Pb or DEHP exposures deflect normal age-related gene expression, DNA methylation programs, and cellular composition across the life course, long after cessation of exposure, and highlight potential biomarkers of developmental toxicant exposures. Further studies are needed to investigate how these epigenetic and transcriptional changes impact cardiovascular health across the life course.

## Introduction

Cardiovascular diseases (CVDs) comprise an array of conditions, including atherosclerosis, heart failure, myocardial infarction, hypertension, cardiac arrhythmias, congenital heart defects, and stroke [1, 2]. In spite of advancements in prevention, diagnosis, and treatment, CVDs remain a leading cause of death in the United States and around the world [3, 4]. CVD risk and pathogenesis are strongly influenced by several factors, including sex, age, diet, lifestyle, and environmental exposures [5–8]. Among these variables, age is the strongest risk factor [9]. Aging is reflected in the both the passage of time since birth (chronological aging), as well as the declines in physiological processes that occur over time (biological aging) [10]. Genetic, environmental, and lifestyle factors may influence the rate of biological aging, resulting in a biological age that is higher or lower than one’s chronological age [10–12]. Notably, several factors that accelerate biological aging are associated with CVDs, including genetic progeroid syndromes, diet, exercise, cigarette smoking, psychosocial stress, and cancer treatments [10, 13–15]. Aging in the cardiovascular system is characterized by widespread changes in the epigenome, including alterations in DNA methylation and chromatin organization [16]. Recent work demonstrates that DNA methylation signatures of accelerated aging are associated with diminished cardiovascular health and increased CVD [17, 18]. However, despite this evidence, several unanswered questions remain. First, it is unclear how exposure to common environmental pollutants during critical windows of development influence the trajectory of age-related cardiovascular epigenetic programming and gene expression across the life course.

Second, although CVDs exhibit marked sexual dimorphism [5, 6, 19], the interplay between sex, developmental pollutant exposure, and aging in the context of the cardiovascular system is poorly understood. The metal lead (Pb) and the plasticizer di(2-ethylhexyl)phthalate (DEHP) are chemically distinct environmental pollutants that differ greatly in their toxicokinetics and toxicodynamics [20, 21]. Nevertheless, both chemicals are strongly associated with various CVDs. In the United States, sources of human exposure to Pb include legacy drinking water systems, contaminated household dust and soil, imported food and consumer products, aviation fuel, and industrial processes [22, 23]. In spite of numerous initiatives to ban the use of Pb in paints, gasoline, and other consumer products worldwide, Pb exposure remains a significant public health threat.

DEHP is a plasticizer widely used in building materials, plumbing, toys, food packaging, pharmaceuticals, and medical tubing [24]. Although the US government and some states have set guidelines regulating DEHP levels in air, water, and consumer products, human exposure to this chemical is still widespread [25]. Both Pb and DEHP exposure are associated with numerous adverse cardiovascular outcomes in human population studies, including congenital heart defects, heart failure, hypertension, cardiac arrhythmias, myocardial infarction, and stroke in Pb exposed individuals [26–30], and hypertension, atherosclerosis, coronary artery disease, decreased heart rate variability, and increased all-cause CVD mortality in individuals exposed to DEHP and its metabolites [31–36].

Developmental exposures to both Pb and DEHP have been shown to impact the epigenome in non-cardiac tissues in human and animal studies, [37–42] and we recently demonstrated that perinatal exposure to these chemicals also impacts sex-specific DNA methylation in the heart [43, 44]. However, the effects of environmental contaminant exposures during early development on age-related epigenetic regulation of gene expression, and potential differences by sex, have not been investigated. To address this knowledge gap, in this study, we examined the effects of exposure to Pb and DEHP during gestation and lactation on the normal trajectory of age-related DNA methylation and gene expression in the heart in both male and female mice.

## Materials and Methods

### Mice and exposure paradigm

The work outlined here was part of a larger developmental exposure study conducted by the National Institute of Environmental Health Sciences (NIEHS) Toxicant Exposures and Responses by Genomic and Epigenomic Regulators of Transcription (TaRGET II) Consortium, which sought to determine how developmental environmental exposures impact the epigenome in multiple tissues and time points across the life course [45]. Mouse Pb and DEHP exposures were performed as outlined previously [44, 46]. Mice utilized for these experiments were wild-type a/a non-agouti mice derived from a colony of the viable yellow agouti (A^vy^) strain maintained in the male line for more than 230 generations. This results in forced heterozygosity on an invariant genetic background and mice that are 93% identical to the C57BL/6J strain [47, 48]. The exposure period began in periconception (2 weeks prior to mating), continued through gestation and lactation, and stopped at weaning at 3 weeks of age. Two weeks prior to mating with virgin a/a males, 8-10 week old virgin females were randomly assigned to control, DEHP (via chow), or Pb (via drinking water) exposure groups. DEHP was dissolved in 7% corn oil and administered via chow (25 mg per kg chow). This results in an estimated maternal DEHP dose of 5 mg/kg/day, assuming that pregnant and lactating female mice weigh approximately 25 g and eat 5 g of chow per day. The level of DEHP exposure falls within the range of human exposure [49], and is within or below the range of no observed adverse effect levels (NOAELs) reported by US and European agencies [50, 51]. Pb II acetate trihydrate (Sigma Aldrich) was dissolved in distilled water to create a 50 mM stock solution, which was then diluted into drinking water at a concentration of 32 ppm. This results in an environmentally relevant maternal blood lead level of 16-60 µg/dL [52]. Pb concentrations in water were verified using inductively coupled mass spectrometry with a limit of detection of 1.0 µg/L (ICPMS; NSF International, Ann Arbor, MI, USA). Exposures were stopped at weaning, and mice from all exposure groups were administered standard chow and drinking water for the duration of the study. The exposure paradigm is outlined in **Figure 1**.

**Figure 1:**
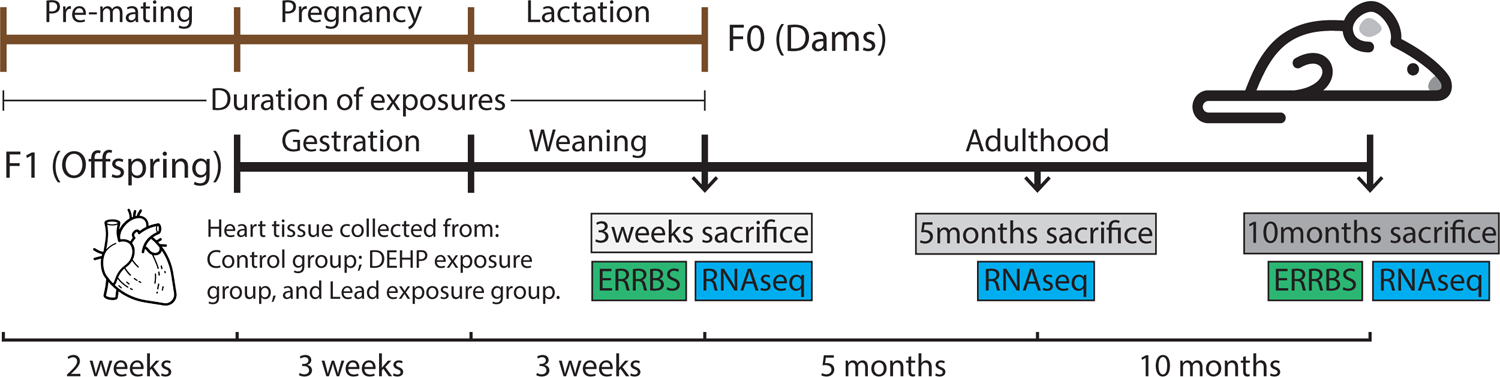
Diagram depicting exposure paradigm and overall experimental design. Dams were exposed to control, DEHP, or Pb beginning 2 weeks prior to mating, and exposure continued through gestation and lactation. Exposures ceased at weaning, and separate cohorts of male and female mice were sacrificed at 3 time points: weaning, 5 months of age, and 10 months of age. ERRBS was conducted in hearts from offspring at 3 weeks and 10 months of age, and RNA-seq was conducted in hearts from all 3 time points.

### Animal monitoring, euthanasia and tissue collection

Animals were monitored daily for signs of illness or distress and were weighed weekly. Approximately 1 male and 1 female offspring per litter were sacrificed at three time points: At weaning on postnatal day 21, at 5 months of age, and at 10 months of age. These time points represent infancy, early adulthood, and middle age, respectively, in humans [53]. Thus, this period spans early childhood and the pubertal transition, and extends through midlife. At sacrifice, a final weight measurement was collected, and mice were euthanized according to protocols established by the NIEHS TaRGET II Consortium [54]. Hearts were extracted, weighed, snap frozen in liquid nitrogen, and stored at −80°C until further processing. RNA and DNA were extracted using the AllPrep kit (Qiagen) according to manufacturer instructions.

### Enhanced reduced representation bisulfite sequencing

Enhanced reduced representation bisulfite sequencing was performed at the University of Michigan Epigenomics and Advanced Genomics Cores as outlined previously [54, 55]. 50 ng of genomic DNA was utilized for each sample. DNA quality was assessed using the Qubit (ThermoFisher, Waltham, MA, USA) and 2200 TapeStation systems (Agilent Technologies, Santa Clara, CA, USA), respectively. Bisulfite conversion efficiencies for all samples were at least 98.4% (average 99.3), and the average mapping efficiency was 65.3%. QC criteria are outlined in Supplementary Table 1. Sequencing was conducted on the Illumina NovaSeq 6000 using an S1 100-cycle flowcell, with an average sequencing depth of ∼49M. On average, this method captured 5% of genomic CpGs.

### RNA-seq

Library preparation was carried out at the University of Michigan Advanced Genomics Core using the KAPA mRNA Hyper Prep Kit (Roche, Wilmington, MA, USA) with dual indexing adapters. Library quality was verified using the Agilent 2200 TapeStation (Agilent, Santa Clara, CA, USA). Sequencing was performed using the Illumina NovaSeq 6000 using an S2 flow cell and paired end, 50bp reads. Average sequencing depth was ∼60M.

### Quality control of ERRBS and RNA-Seq data

For quality control of ERRBS and RNA-Seq data, FastQC (v0.11.8) was used to assess the overall quality of each sequenced sample. TrimGalore (v0.4.5) was applied to removed adaptor sequences and trim low-quality bases. After trimming, reads with length less than 20 nucleotides were removed from further analysis for both ERRBS and RNA-Seq. Bismark (v0.22.1) with Bowtie2 (v2.3.4) as backend was used for reads alignment and methylation calling for the ERRBS data with default settings (multi-seed length of 20bp with 0 mismatches). The unmethylated lambda phage DNA was used to calculate the bisulfite conversion rates.

STAR (v2.7.1a) was used to perform reads alignment for RNA-Seq data. HtSeq-count (v0.11.2) with Python (v3.7.3) was applied to generate the final reads count. Genome Reference Consortium Mouse Build 38 (mm10) was used as the reference genome for both ERRBS and RNA-Seq data.

### DEG and DMR Analysis

The Bioconductor packages RUVSeq (v1.32.0) and edgeR (v3.40.2) were used to correct the batch effects and identify the differentially expressed genes, respectively. The RUVr function with *k*=3 was applied to remove any potential batch effects among the three time points of the RNA-Seq data within each treatment group. Genes with at least 5 reads counted in at least 12 animals were kept for further DEG analysis. To identify aging related DEGs, three comparisons, i.e. PND21 vs. 5 Month, 5 Month vs. 10 Month, and PND21 vs. 10 Month, were performed within each treatment group. Male data and female data were analyzed separately. Significant DEGs were defined for all comparisons as those genes with FDR less than 0.05 and absolute log fold change greater than 2.

The Bioconductor package methylSig (v0.5.2) was used to identify differentially methylated regions. A tiling window with 100 nucleotides was applied for DMR detection. CpG sites with less than 10 and more than 500 reads covered were removed from further analysis. Tiling windows with required coverage in at least four samples per comparison group were used for DMR detection. DMRs were identified using the *methylSigDSS* function between PND21 and 10 month within each treatment group and sex. An FDR less than 0.05 and methylation change larger than 10% were used to select significant DMRs. The Bioconductor package annotatr (v1.24.0) was used to annotate the DMRs to different genomic regions, including CpG islands, CpG shores, CpG shelves, CpG intervals, promoters, exons, introns, 5’UTRs, 3’UTRs, enhancers, and 1-5 kb upstream of TSSs.

### Gene Ontology Analysis of DEGs and DMRs

The Bioconductor packages clusterProfiler (v4.6.2) and ChipEnrich (2.22.0) were used to perform Gene Ontology (GO) analysis with DEGs and DMRs, respectively. The *enrichGO* function in clusterProfiler with DEGs as input was used to identify related GO terms. For DMRs, the *chipenrich* function with locus definition *nearest_tss* (the region spanning the midpoints between the TSSs of adjacent genes) was applied to discover enriched GO terms. All three ontologies, Biological Process (BP), Cellular Component (CC), and Molecular Function (MF), were used in both DEG and DMR related GO analysis. An FDR < 0.05 cutoff was used to select significantly enriched GO terms.

### Cell type deconvolution of bulk RNA-seq data

To quantify whether changes with exposure or aging were associated with alterations in cell type composition of heart tissues, we used a bioinformatic deconvolution method based on a single cell atlas of the normal heart, based on data generated as part of the Human Cell Atlas [56]. Sample-specific counts matrices of single cell RNA-seq profiling of hearts were downloaded from the Human Cell Atlas Data Explorer and loaded into Seurat [57]. These data were downsampled to 25,000 cells via the “subset” function and we then used these single cell gene expression data to predict the cellular composition of our tissues based on their bulk RNA-seq profiles. For this deconvolution, we used the Multi-subject Single Cell (MuSiC) deconvolution method [58], which predicts cell type proportions in bulk RNA-seq data based on a reference multisubject single cell RNA-seq dataset. Mouse-to-human gene alignment occurred with BioMart [59], and then MuSiC uses a non-negative least squares regression-based method based on the cell type specific gene expression signatures from the single cell data, with constraints that individual cell type proportions must be a positive value and their sum cannot exceed 1. The cell proportion differences over time or exposure were tested for statistical significance by one-way analysis of variance (ANOVA).

## Results

### Age-related changes in DNA methylation in control and toxicant exposed animals

To examine how Pb and DEHP exposure affected age-related DNA methylation, we conducted ERRBS in whole hearts at weaning and 10 months of age (**Figure 2A**), with each treatment group having 5-7 animals per sex (**Figure 2B**). We determined the number of age-related changes in DNA methylation (i.e. between weaning and 10 months of age) in each treatment group, which is summarized in **Figure 2C**. In each group, we observed several thousand age-related DMRs, with the majority (78%-84%) being hypermethylated (**Figure 2C**). In both sexes, Pb and DEHP exposure resulted in a slightly greater percentage of DMRs showing hypermethylation compared to control (**Figure 2C**). We next annotated the DMRs in each condition to the mouse mm10 genome and found that enrichment of age-related DMRs differed based on the direction of methylation change. Compared to all genomic regions tested, hypomethylated DMRs were less likely to be found in CpG islands, shelves, exons, introns, 3’UTRs, and enhancers. In contrast, hypermethylated DMRs were slightly enriched for several of these regions, including CpG islands, exons, introns, and enhancers (**Figure 2D**). These patterns were qualitatively similar across sex and exposure group. Lists of annotated, age-related DMRs for each sex and exposure group are included in Supplementary Table 2.

**Figure 2:**
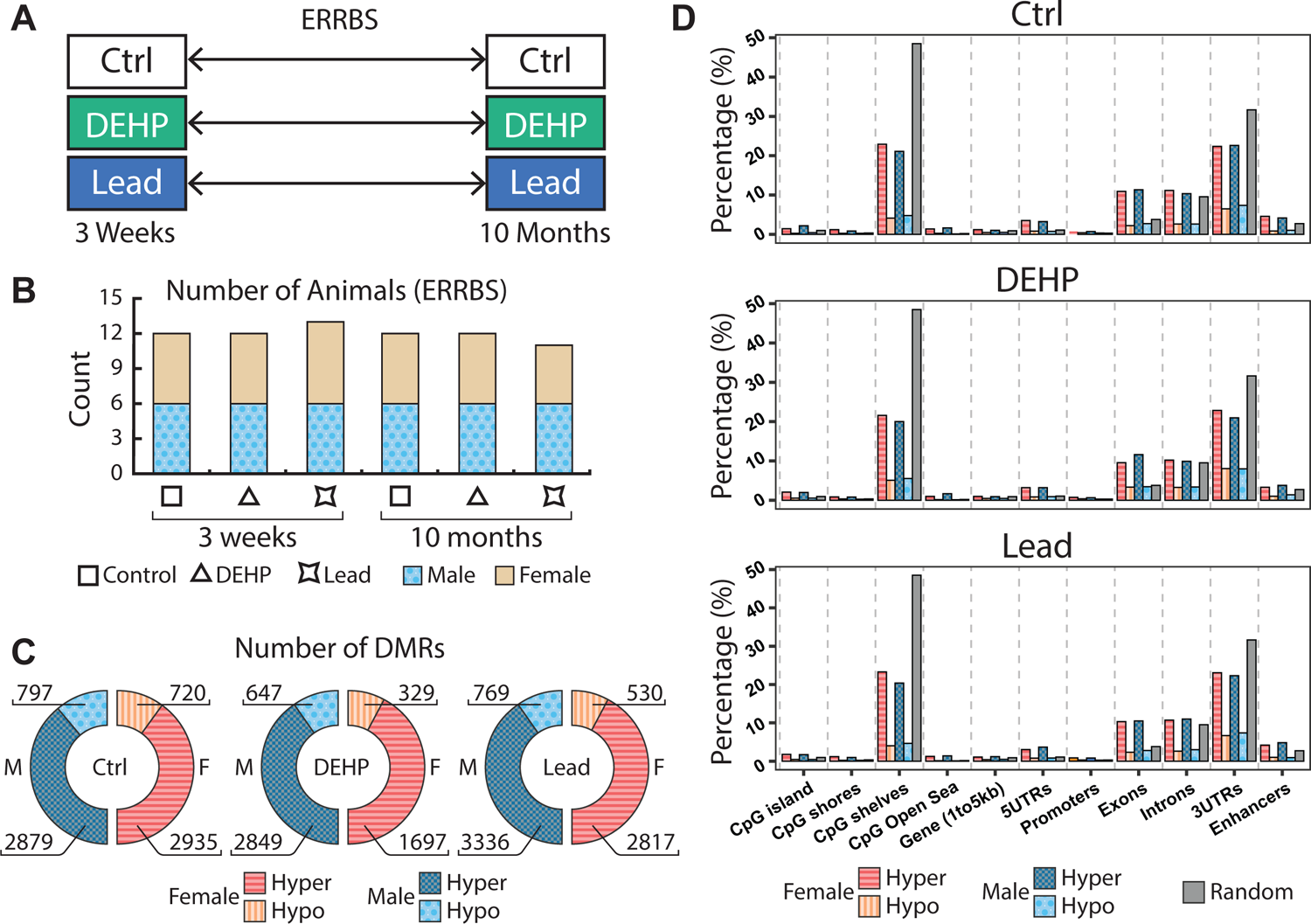
ERRBS in offspring mouse hearts. A. Age-related changes in cardiac DNAm were determined by comparing 3 week hearts with those of animals at 10 months of age. This comparison was conducted within each exposure group (control, DEHP and Pb). B. Number of animals in each sex/exposure group. C. Number of age-related DMRs in each exposure group, with hypomethylated DMRs depicted in lighter shading, and hypermethylated DMRs in darker shading. D. Percentage of DMRs mapping to various regions of the genome, compared to what would be expected in a random distribution, stratified by sex and direction of methylation change.

### Age-associated DNA methylation differs by sex and exposure

In order to understand the biological pathways undergoing differential DNA methylation, we conducted pathway analysis using ChIP-Enrich, stratifying by sex, exposure, and direction of methylation change. The results of this analysis are shown in **Figure 3** and in Supplementary Table 3. **Figure 3A** summarizes significant Gene Ontology Biological Process (GOBP) terms for each combination of sex, exposure, and direction of methylation change which are relevant to cardiovascular development and disease. An upset plot depicting overlaps among all of the enriched GO terms (GOBP, GOCC, GOMF) is shown in Supplementary Figure S1, and depicts that the majority of enriched GO terms fall within a single condition. Overall, differential DNA methylation occurred in pathways associated with differentiation, disease, and development (**Figure 3A** and Supplementary Table 3). A closer examination of the significant pathways specifically related to cardiovascular development and disease showed that the majority of enriched pathways occurred within a single sex/exposure combination, although pathways related to embryonic development, pattern specification, response to vascular endothelial growth factor stimulus, animal organ regeneration, and cardiac hypertrophy were enriched in were enriched in multiple conditions (**Figure 3A**). Enriched pathways also differed based on the direction of DNA methylation change, underscoring the importance of stratifying on the basis of hyper vs. hypermethylated DMRs. Notably, a few pathways were enriched in exposed animals but not in control or vice-versa. For example, the hypomethylated DMRs in DEHP exposed males and females were enriched for cardiac muscle hypertrophy, while hypo and hypermethylated control females, but not exposed animals, showed enrichment for animal organ regeneration (**Figure 3A**). **Figure 3B-E** illustrates the total number of significantly enriched GOBP pathways for each condition. The numerator of each fraction represents the number of pathways containing heart-specific terms (muscle, ventricular, atrial, cardiac, aorta, and heart), and the denominator all other pathways, and the data show that the number of total and heart-specific pathways differ across exposure groups. These results collectively demonstrate that age is associated with altered DNA methylation in pathways related to development and disease, with marked differences based on sex and exposure.

**Figure 3:**
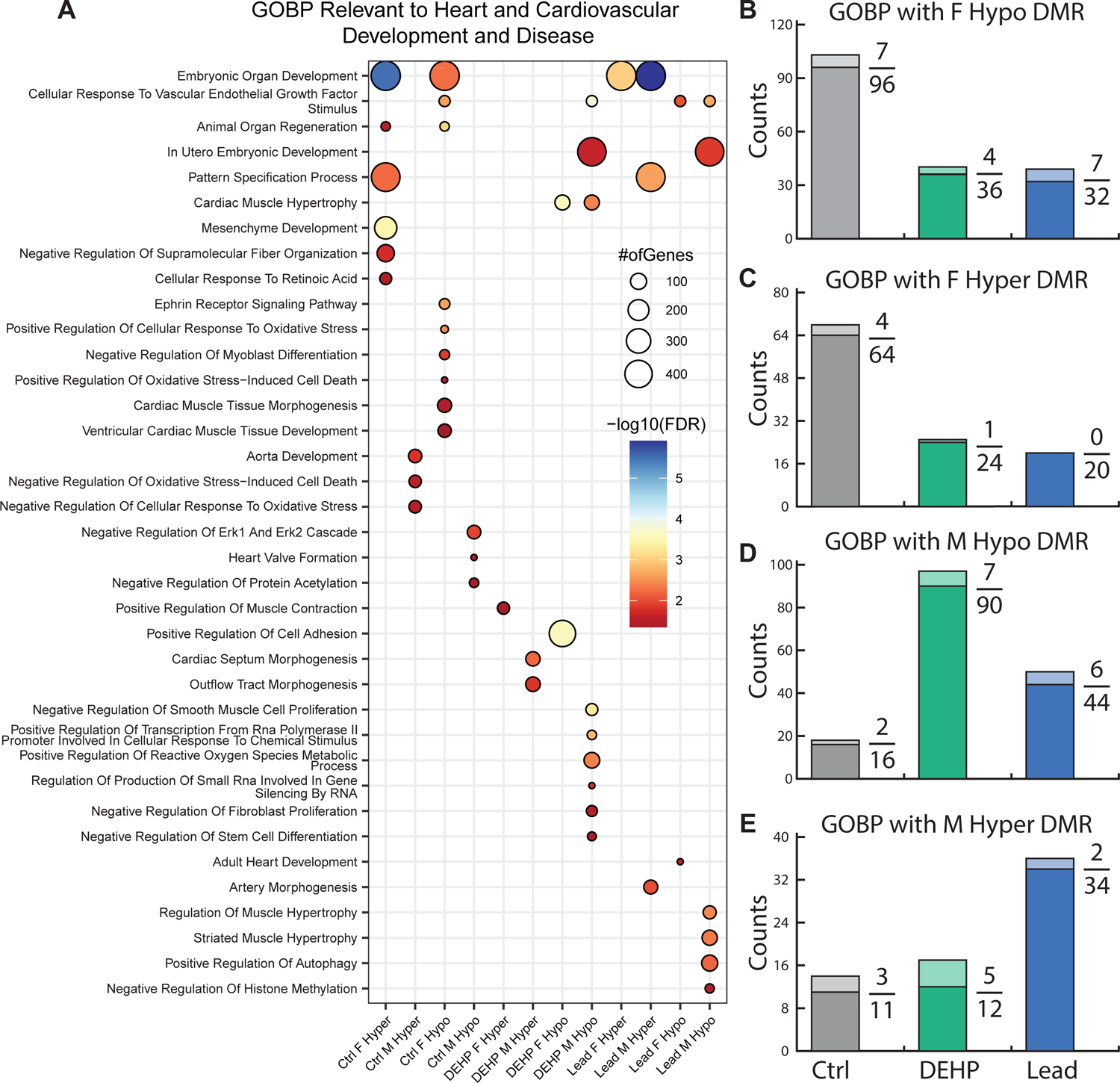
Pathway analysis of age-related changes in DNAm. A. Summary of the significant Gene Ontology Biological Process (GOBP) terms for each combination of sex, exposure, and direction of methylation change which are relevant to cardiovascular development and disease. The size of each circle reflects the number of genes in each category, and color indicates the -log10(FDR). B.-E. Number of enriched GOBP pathways for each combination of sex, exposure, and direction of methylation change. The numerator of each fraction depicts the number of pathways containing heart-related terms (muscle, ventricular, atrial cardiac, aorta, and heart) and the denominator depicts the number of all other pathways.

### Age-related changes in gene expression in control and exposed animals

We next examined how the cardiac transcriptome changed with age in males vs. females, and the effects of chemical exposures on this process. To this end, we conducted RNA-seq on samples of RNA from the same hearts utilized for ERRBS (5-7 animals per condition, **Figure 4A-B**) at weaning and 10 months of age, as well as an additional cohort of animals at 5 months of age. Volcano plots depict the number, magnitude, and direction of changes in gene expression for each sex and exposure, comparing weaning and 10 months of age (**Figure 4C-H**). Volcano plots depicting comparisons between weaning vs. 5 months of age, and 5 months vs. 10 months of age, are shown in Supplementary Figure S2. Numbers of differentially expressed genes (DEGs) are also summarized in Table 1, and the full lists of DEGs can be found in Supplementary Tables 4-6. We found several hundred DEGs between weaning and 10 months of age in all exposures/sexes, with the largest number of DEGs (933) occurring in control female hearts (Table 1). Within each exposure group, the majority of age-related changes in gene expression were sex specific; however, there were several genes in common across sexes (**Figure 5A**). When we compared age-related gene expression changes across exposure groups, we found that the majority of differentially expressed genes were unique to each exposure (**Figure 5B**). This finding suggests that DEHP and Pb exposure considerably altered the normal age-related gene expression profile in the heart across the life course. To explore this further, we conducted RNA-enrich pathway analysis to assess the gene pathways differentially expressed between weaning and 10 months of age in each group. Full lists of enriched GO pathways can be found in Supplementary Tables 7-9. **Figure 5C** illustrates the number of significantly enriched GOBP pathways for each group, as well as the number of pathways overlapping multiple conditions. This analysis revealed little overlap in differentially expressed pathways across the different conditions. A single pathway (extracellular matrix organization) was significantly altered in all sexes and exposure groups, while Pb exposed males and females had the largest number of overlapping pathways (11). Two pathways (tissue remodeling and muscle contraction) were significantly differentially expressed in all exposed groups but not in control, and there was 1 pathway unique to both male and female control groups (protein kinase B signaling). These data further support the hypothesis that gene expression changes across the life course are deflected by toxicant exposures during early development.

**Figure 4:**
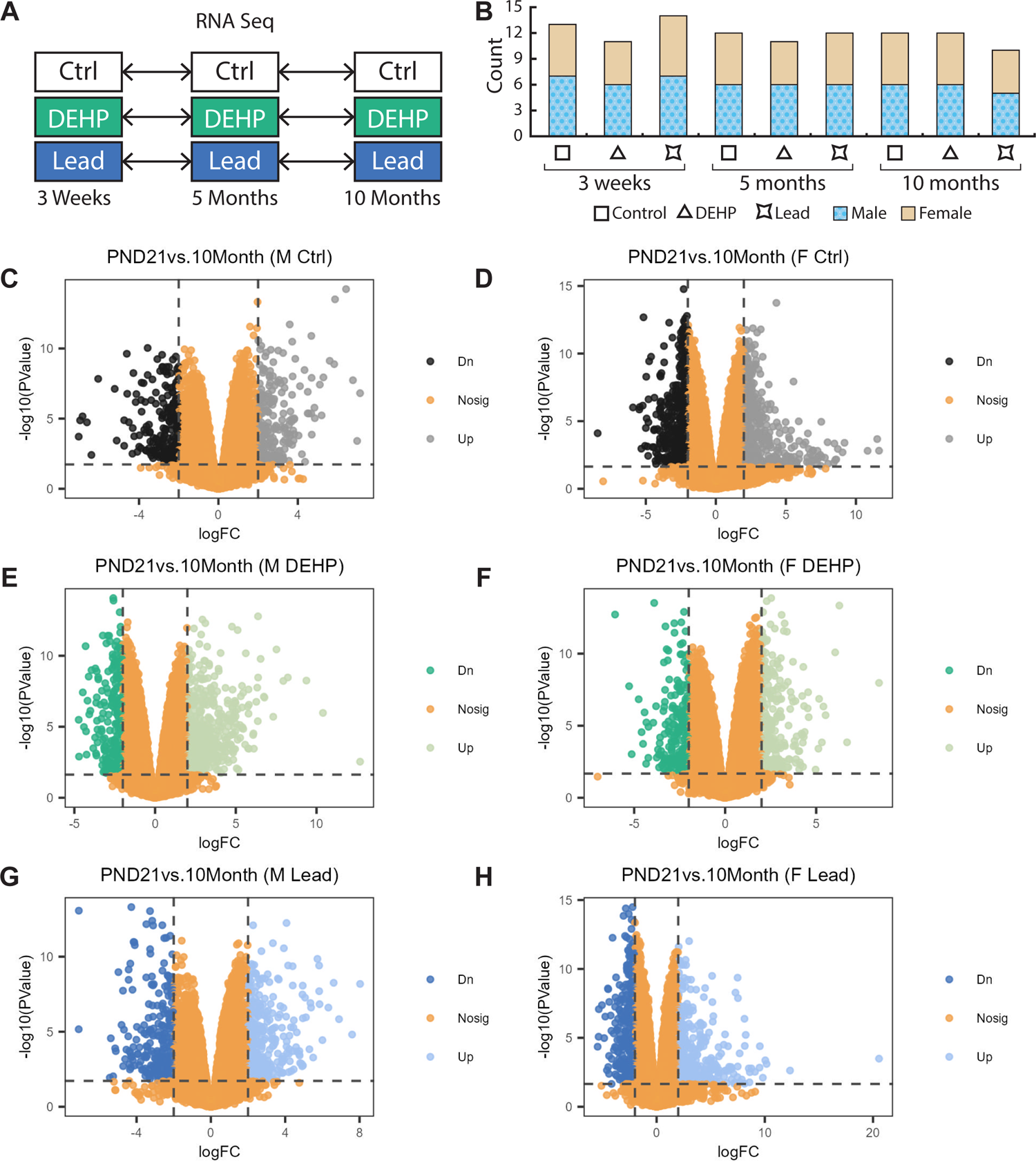
Effects of age and exposures on gene expression. A. Age-related changes in cardiac gene expression were determined by comparing weanling hearts with those of animals at 5 months and 10 months of age within each exposure group. B. Number of samples in each sex and exposure group. C-H. Volcano plots showing the number of significantly up or down-regulated genes between weaning and 10 months of age. Differentially expressed genes were defined as those with a FDR<0.05 and an absolute logFC >2.

**Figure 5:**
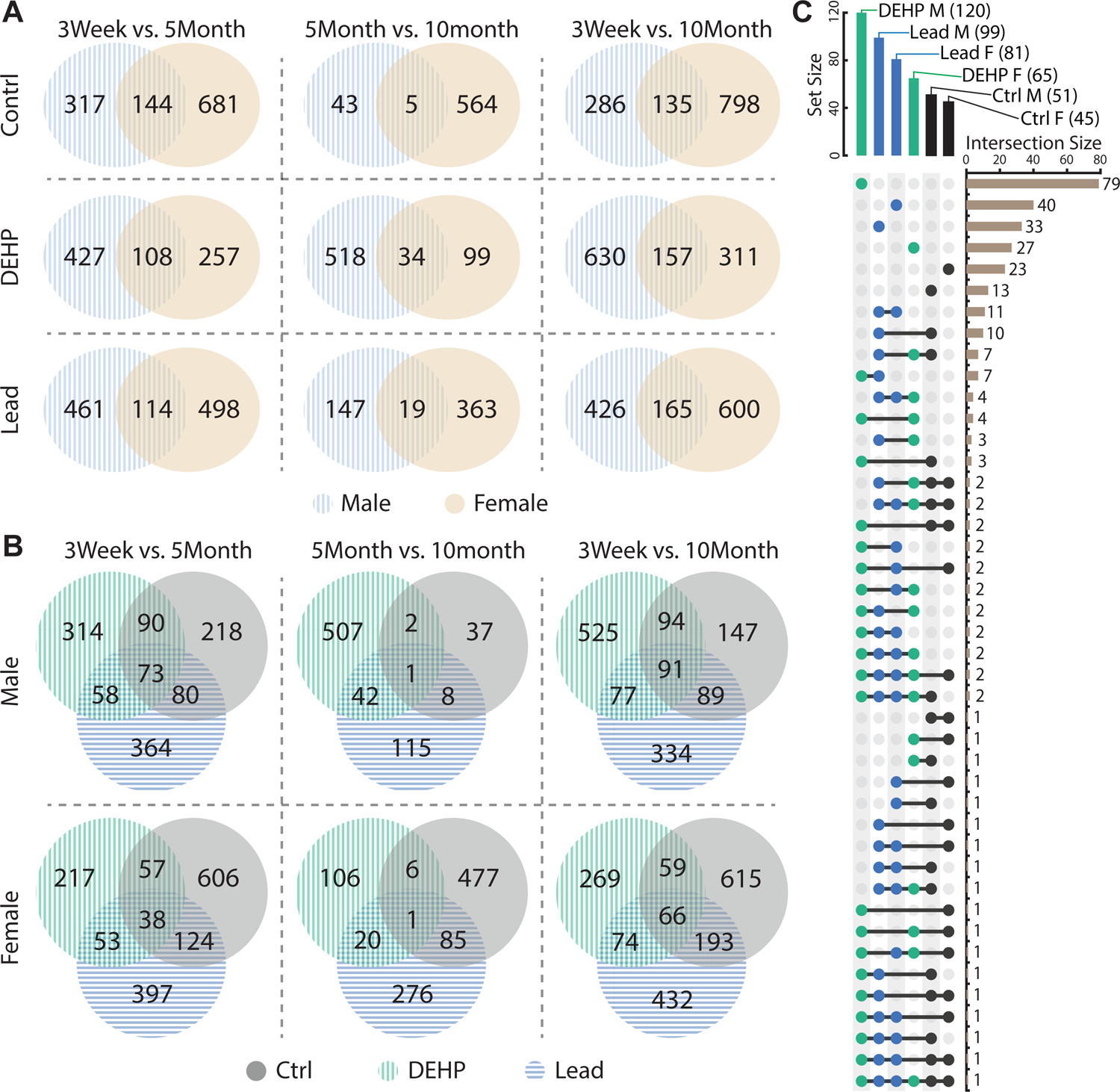
Sex and exposure specificity of age-related transcriptomic changes. A. Venn diagrams showing sex overlap in differential gene expression for each time bracket comparison and exposure. B. Venn diagram showing exposure overlap in differential gene expression for each time bracket comparison and sex. C. UpSet plot showing the number of intersecting GOBP pathways for each exposure and sex combination.

**Table 1:** Number of differentially expressed genes (DEGs) between 3 weeks and 5 months of age, between 5 months and 10 months of age, and between 3 weeks and 10 months of age. Significance was based on an FDR less than 0.05 and absolute log2(fold change) > 2.

| Exposure | 3 weeks vs. 5 months |  | 5 months vs. 10 months |  | 3 weeks vs. 10 months |  |
| --- | --- | --- | --- | --- | --- | --- |
|  | Male | Female | Male | Female | Male | Female |
| Control | 461 | 825 | 48 | 569 | 421 | 933 |
| DEHP | 535 | 365 | 552 | 133 | 787 | 468 |
| Pb | 575 | 612 | 166 | 382 | 591 | 765 |

### Concordant age-related changes in gene expression and DNA methylation

We next examined the extent to which the observed age-related changes in DNA methylation and gene expression in control and exposed animals occurred at the same genes. To this end, we identified all age-related DEGs occurring between any of the 3 time point comparisons (weaning vs. 5 months, 5 months vs. 10 months, and weaning vs. 10 months) and determined whether they also showed differential DNA methylation between weaning and 10 months of age. As shown in **Figure 6A**, several genes in each group showed concomitant changes in DNA methylation and gene expression. Full lists of these genes, as well as separate Venn diagrams for each of the 3 time point comparisons, can be found in Supplementary Table 10 and Supplementary Figure S3. Notably, a subset of genes showed significant age-related differential methylation and expression in exposed but not control hearts (25 and 20 genes in males and females, respectively). Several of these genes exhibited age-related expression trajectories that differed by sex and exposure, including *Krt18* (found in DEHP male, Pb male, and Pb female), *Atp8a2* (found in DEHP and Pb-exposed females), *Ston2* (found in DEHP and Pb-exposed females, Pb male), and *Pou3f1* (found in DEHP and Pb-exposed males) (**Figure 6B-E**).

**Figure 6:**
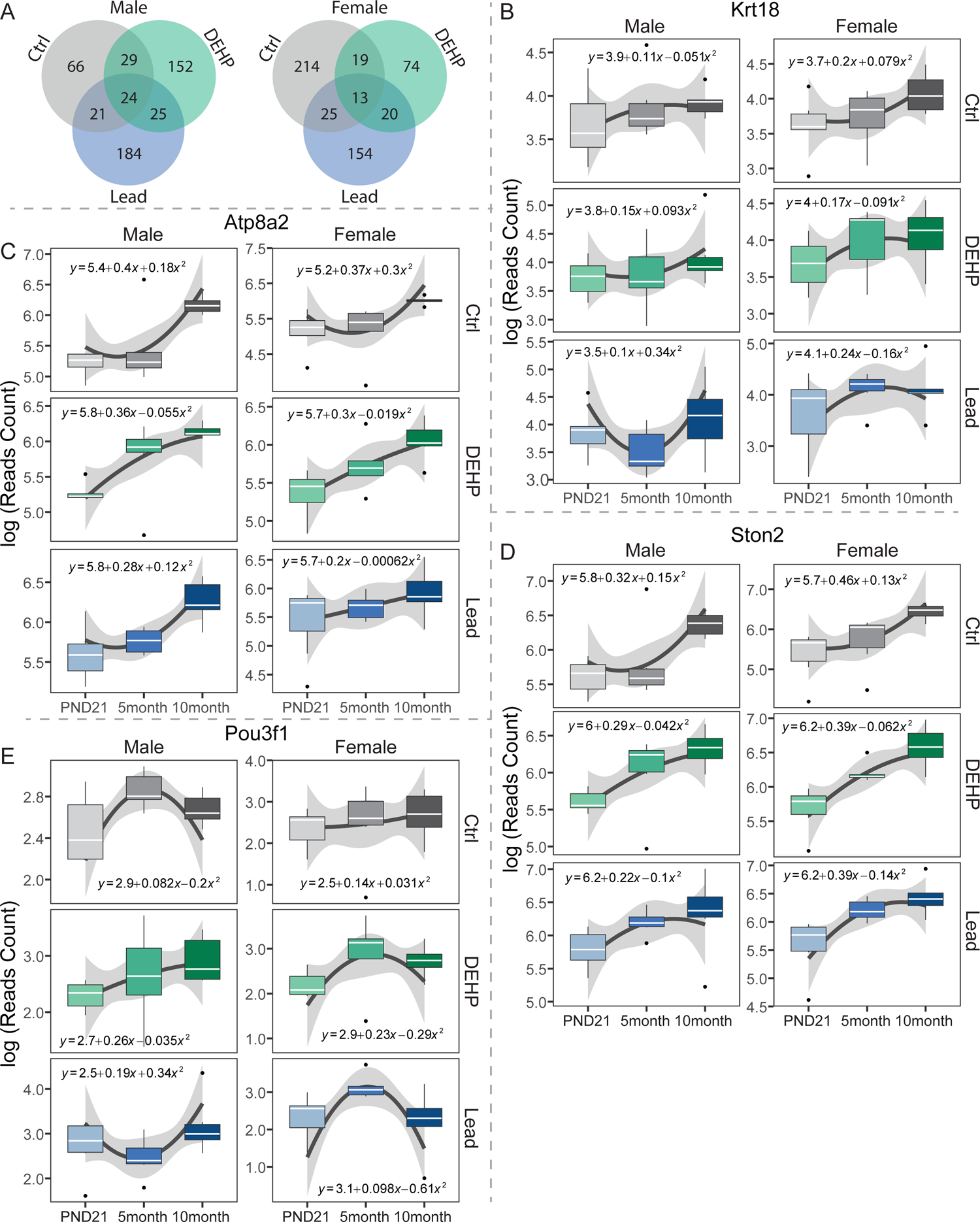
Genes with differential expression and DNA methylation. A. Venn diagrams showing the number of genes in each sex exhibiting differential expression in any of the 3 time point comparisons (weaning vs. 5 months of age, 5 months of age vs. 10 months of age, and weaning vs. 10 months of age) and differential methylation B.-D. RNA-seq read count data for 4 genes showing that the age-related gene expression trajectory differs by exposure and/or sex. Each of the 4 genes were differentially expressed and methylated with one or both exposures but not in control.

### Age-related shifts in cell type

The heart is comprised of several different cell types, including cardiomyocytes, fibroblasts, endothelial cells, pericytes, smooth muscle cells, adipocytes, immune cells, and neuronal and glial cells [56]. Shifts in cardiac cellularity across the life course have been reported, including increased fibrosis in old age [60] as well as changes in the composition of epicardial fat tissue [61], but the effects of developmental exposures on age-related changes in cellularity between weaning and middle adulthood are unknown. Therefore, we conducted cell type deconvolution of the RNA-seq data using a published algorithm and publicly available single cell transcriptomic data [56, 58] and quantified the relative proportions of atrial and ventricular cardiomyocytes, endothelial cells, smooth muscle cells, fibroblasts, adipocytes, lymphoid and myeloid cells, neuronal cells, pericytes, and mesothelial cells. Endothelial and smooth muscle cells were present in the largest proportions, followed by atrial and ventricular cardiomyocytes, while neuronal and mesothelial cells comprised the smallest proportions of cells (**Figure 7**). Summary statistics for all of the comparisons by age and exposure can be found in Supplementary Table 11. Overall, age-related trends in cell composition were generally consistent across sexes and exposure groups. The proportions of atrial cardiomyocytes remained consistent with age in all conditions (**Figure 7A**). Proportions of ventricular cardiomyocytes increased between weaning and 10 months of age across all conditions (p>0.05, **Figure 7B**). However, in males exposed to Pb and DEHP, the increase in ventricular cardiomyocytes began at 5 months of age, in contrast to controls which did not show differences until 10 months of age (**Figure 7B**). In mice from both sexes, there was an age-related decline in the proportion of smooth muscle cells in control and DEHP-exposed mice (p<0.05), but no change with age in Pb-exposed mice (**Figure 7C**). Age-related trends in the proportions of endothelial cells did not differ across exposure groups (**Figure 7D**). In control and exposed males, fibroblasts significantly decreased between weaning and 10 months of age (p>0.05), a trend that was only present in Pb-exposed females (**Figure 7E**). The proportions of adipocytes increased across time in all conditions (p<0.05), with no differences by exposure (**Figure 7F**). Pericyte proportions significantly increased with age in control males (p<0.05), but this change was not observed in exposed males or in females under any condition (**Figure 7G**). Proportions of neuronal cells did not significantly differ with age, except in Pb-treated males and females (p<0.05), where they decreased from a higher baseline between weaning and 10 months of age (**Figure 7H**). Lymphoid cell proportions trended up with age, a pattern that reached statistical significance in Pb-exposed females as well as control and DEHP exposed males (**Figure 7I**). Myeloid cells were significantly higher at 10 months of age compared to weaning in all conditions except Pb-exposed females, which exhibited a trend toward significance (**Figure 7J**). No age-related changes in mesothelial cells were observed in any of the conditions (**Figure 7K**). Collectively, these results demonstrate that there are small but significant shifts in cell proportions with age that, in some cell types, are influenced by sex and/or exposure.

**Figure 7:**
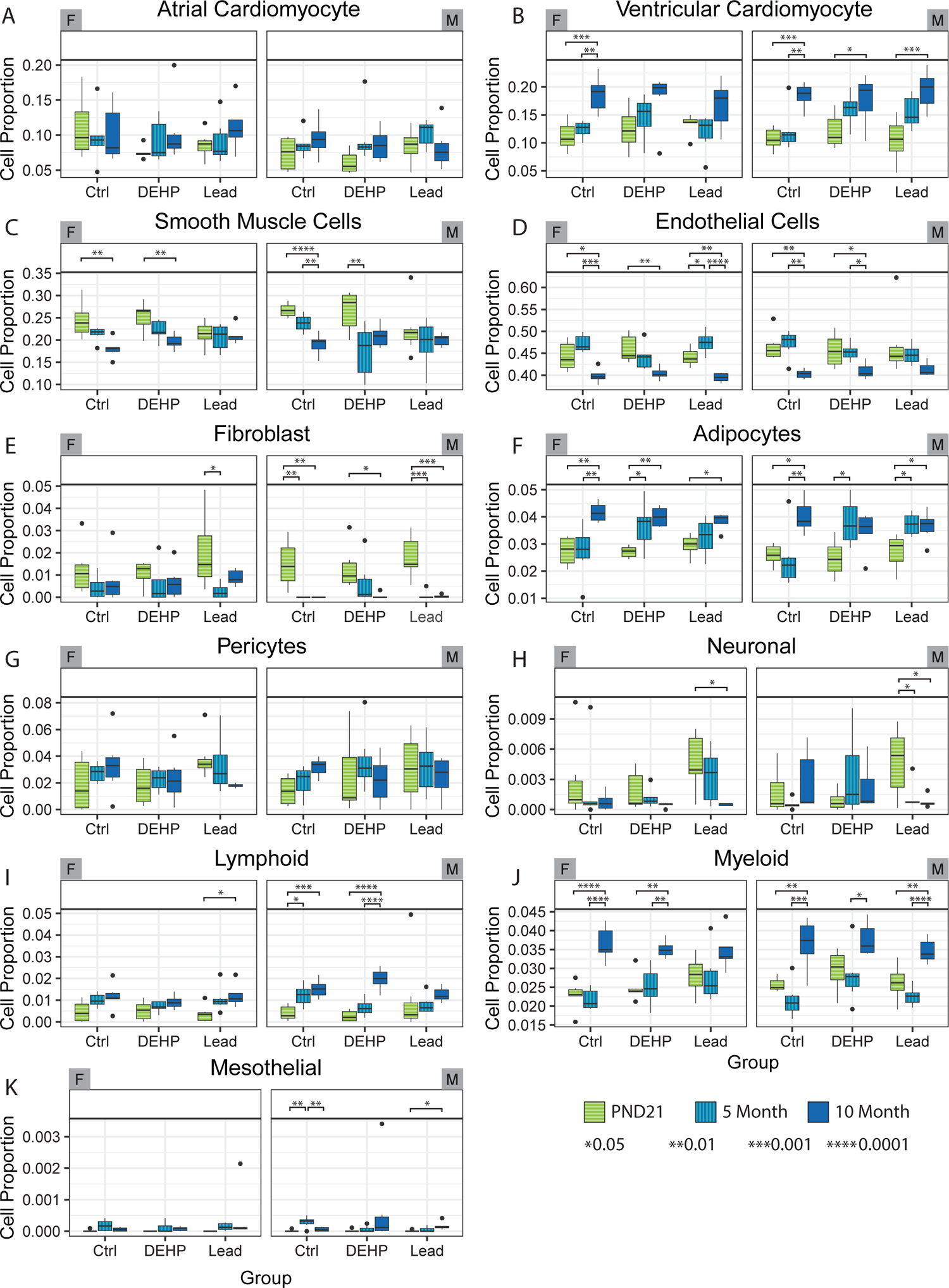
Cell type deconvolution analysis of whole heart tissue, showing the relative proportions of the various cell types found in male and female hearts in each time point and exposure group. P-values depict statistically significant differences in cell proportions across time within each exposure group (one way ANOVA with pairwise t-test). Comparisons across exposure within each age group were also conducted and can be found in Supplementary Table 11.

## Discussion

Cardiac development begins in early embryogenesis, and growth and maturation continue during the postnatal period and adolescence [62]. Although development is largely complete by adolescence, the heart continues to undergo normal age-related changes in physiology, at a rate which varies based on the individual [63]. Cardiac development and aging are characterized by widespread transcriptional, epigenetic, and metabolic changes [64–66], but how chemical exposures may interfere with these processes, and the potential health effects, are unknown. Studies showing widely divergent age-related epigenetic patterns in pairs of twins underscore the influence of environmental factors on the aging epigenome [67–69]. Such environmental deflection of the normal aging process, as we have previously defined it [70], may have important implications for long-term disease risk.

In this study, we demonstrate that developmental exposure to two chemically distinct and ubiquitous environmental contaminants causes marked deflection of the normal age-related trajectory of DNA methylation and gene expression occurring between weaning and middle age. Multiple age-related epigenetic changes have been reported [66, 71], with the majority of studies thus far focusing on DNA methylation. In humans, aging is associated with altered DNA methylation at various stages across the life course, including in childhood [67, 72], between early childhood and adolescence [73, 74], and during adulthood and old age [75, 76]. Age-related changes in DNA methylation have been characterized in numerous human tissues, including peripheral blood and various blood cell types, buccal epithelium, brain, kidney, skeletal muscle, prostate, liver, adipose, and cervix [67, 76–79], and similar profiling studies have been conducted in rodent tissues [80–82]. To date, no investigations into the effects of aging on DNA methylation and gene expression in the heart have been conducted in humans. Although one study did profile age-related changes in DNA methylation the mouse heart, females were not included [81]. Collectively, the aforementioned human and rodent studies published to date show marked tissue specificity in age-related DNA methylation. Indeed, tissue of origin is the most important discriminating factor among DNA methylation samples, even if they come from the same individual [83]. These observations all underscore the importance of profiling the tissue itself, rather than a surrogate tissue, to gain the greatest biological insight into the effects of aging. This study is therefore impactful, as we examine how age affects the epigenome and gene expression in the heart in both males and females, at baseline and in response to two toxicants of high relevance to humans.

Although epigenetic aging studies in twins or non-twin family members [68, 69, 84] strongly suggest that environmental factors impact the aging process, the influence of specific chemicals on epigenetic aging is poorly understood. However, a number of recent studies have begun to shed light on this. In humans, these studies are largely restricted to blood, with few studies reporting effects on tissues targeted by the chemical exposures. Benzene, trichloroethylene, organochlorine pesticides, and PFAS, as well as air pollutants such as tobacco smoke, particulate matter, sulfate, and ammonium have all been associated with accelerated aging as measured via well-established DNA methylation clocks in blood [85–89]. Investigation of human liver samples showed that alcohol dependence is associated with increased DNA methylation age in the liver [90]. Animal studies have provided additional evidence that bisphenol A, Pb, and trichloroethylene can impact epigenetic aging of various tissues, through DNA methylation and other epigenetic factors [91–94]. Mouse studies of Pb exposure and epigenetic aging to date have examined age-related DNA methylation of several loci in tail tissue [92] or expression of specific miRNAs in the brain [94]. As effects of chemical exposures on age-related epigenetic changes in the heart have not been investigated, our study addresses an important knowledge gap in this area.

In this study, we observed that the majority of age-related DMRs are hypermethylated, irrespective of age, sex, or exposure. This finding is in keeping with our previous work, which showed a general trend toward DNA hypermethylation in longitudinal mouse blood samples between 2 and 10 months of age [95]. In contrast, numerous studies demonstrate that aging is associated with genome-wide hypomethylation [96–98]. However, the period between weaning and 10 months of age analyzed in our study encompasses the early perinatal phase, puberty/adolescence, and adulthood. Effects of developmental exposure to these chemicals on old age will be an important area for future investigation. Previously published work in human blood has shown that global DNA methylation increases during early postnatal life, remains stable during adulthood, and then only declines with old age [67, 73, 96, 97]. We also observed hypermethylation of CpG islands and promoters with age in both sexes and in all 3 exposure groups, in accordance with previous work [96–98]. Thus, our findings are consistent with age-related trends observed in other studies. We found that age-related DNA methylation occurred to a large extent in pathways related to tissue development and cell fate commitment. This aligns with work from others showing that age-related DNA methylation occurs at pathways related to development, obesity, longevity, and cancer across multiple species [99]. Although we observed changes in pathways related to development and differentiation across all conditions in general, the specific pathways enriched within each sex and exposure group were largely unique, suggesting that age-related changes in cardiac DNA methylation exhibit sex specificity in unexposed animals, and are altered in distinct ways by developmental exposure to Pb or DEHP. Our transcriptional analysis revealed similar sex-specific changes in gene expression with age, which were deflected by developmental exposure to Pb or DEHP. Notably, pathway analysis of these data showed that Gene Ontology pathways related to Tissue Remodeling and Muscle Contraction were significantly altered between weaning and 10 months of age in males and females exposed to Pb or DEHP. As defects in both processes are implicated in a variety of CVDs [100, 101], it will be important to investigate the long-term effects of Pb exposure on cardiac function, an important future direction of this work.

Although the majority of age-related differential DNA methylation and gene expression did not occur in the same genes, we observed several genes that were both differentially methylated and expressed with age in each exposure group. Several genes, including *Atp8a2*, *Krt18*, *Pou3f1*, and *Ston2* showed age-related gene expression trajectories that differed by sex and/or exposure. Although cardiovascular functions for *Atp8a2* and *Ston2* have not yet been reported, the *Krt18* gene was recently reported to play a protective role in heart failure [102].

*Pou3f1* is a neurogenic factor which has been shown to be upregulated upon epigenetic dysregulation of normal cardiac differentiation [103]. Interrogation of the Comparative Toxicogenomics Database [104] revealed that all four genes are differentially expressed and/or methylated in response to numerous chemicals with diverse mechanisms of toxicity, including bisphenol A and benzo(a)pyrene, suggesting that these genes may represent biomarkers of chemical exposure.

Age- and exposure-related epigenetic changes in whole tissue samples may be due to cell intrinsic changes, but they may also reflect changes in cell type composition [105]. Because cellular composition of the heart is known be influenced by age, disease, and sex [60, 61, 106], we utilized cell type deconvolution to examine the extent to which the observed age-related changes in DNA methylation and gene expression were due to changes in cellular composition. We observed subtle but statistically significant changes in cell type proportions based on age, sex, and exposure. Thus, the age-related changes in gene expression and DNA methylation observed in this study are likely due to both cell intrinsic transcriptional and epigenetic changes, as well as changes in cellular composition in the heart. Future studies using single cell approaches will shed further light on this question.

The health implications of Pb or DEHP mediated deflection of DNA methylation and gene expression in the heart are unclear, but several lines of evidence suggest that age-related changes in DNA methylation may impact long-term cardiovascular health. In mice, age-related DNA methylation is associated with more severe injury following an ischemia-reperfusion event [107]. Several human studies have also linked epigenetic age acceleration with an increased risk of various markers of cardiovascular disease in both white and black populations [17, 108–110], though sex differences and contributions of environmental factors are unclear.

The molecular mechanisms underlying the deflection of age-related DNA methylation by Pb or DEHP exposure are currently unclear. Methylation of DNA is carried out by DNA methyltransferases (DNMT1, DNMT3A, DNMT3B), using S-adenosylmethionine (SAM) as a cofactor. DNA hydroxymethylation, the first step in the process of active DNA de-methylation, is catalyzed by TET dioxygenases (TET1, TET2, TET3), with alpha ketoglutarate (α-KG), iron, and vitamin C as cofactors. Changes in DNA methylation may thus occur though altered expression or function of these enzymes, or depletion of their cofactors. Studies in various cellular and animal models show that both DEHP and Pb can alter expression of DNMTs and/or TETs [111–115]. There is some evidence that both Pb and DEHP may alter levels of SAM, but there is otherwise little known about how these toxicants impact other cofactors for epigenetic modifying enzymes [116, 117]. Pb and DEHP both have been shown to cause oxidative stress in multiple contexts [118, 119] and given that DNA methylation is sensitive to cellular redox status [120], this is another plausible mechanism of epigenetic programming by these chemicals.

This study has a few key limitations. First, these data represent a relatively small sample size in one mouse strain, and further experiments will be necessary to confirm our findings. In addition, to measure DNA methylation, we utilized a sodium bisulfite conversion-based method, which does not discriminate between 5-methylcytosine and other, more oxidized modifications such as 5-hydroxymethylcytosine [121]. Thus, the differentially methylated cytosines and regions reported here reflect a combination of all of these modifications. Although 5-hydroxymethylcytosine is approximately an order of magnitude less abundant than 5-methylcytosine [122], recent studies suggest that the modification has its own distinct molecular functions and plays an important role in cardiac development and disease [123]. Identifying age and exposure-induced changes in 5-hydroxymethylcytosine is an important future direction of this research.

## Conclusion

In summary, we demonstrate herein that developmental exposure to Pb or DEHP shapes the trajectory of age-related DNA methylation and gene expression in the heart in a sex-specific manner, long after cessation of exposure. Given that the risk of cardiovascular disease increases markedly with age, future studies should investigate how these changes impact long-term cardiovascular health.

## Acknowledgements

Funding for this work was provided by the National Institute of Environmental Health Sciences (NIEHS) TaRGET II Consortium U01 (ES026697), a NIEHS Transition to Independent Environmental Health Researcher (TIEHR) K01 (ES032048), a NIEHS Revolutionizing Innovative, Visionary Environmental Health Research (RIVER) R35 (ES031686), a National Institute of Aging R01 (AG072396), the University of Michigan NIEHS/EPA Children’s Environmental Health and Disease Prevention Center P01 (ES022844/RD83543601), the Michigan Center on Lifestage Environmental Exposures and Disease (M-LEEaD), NIEHS T32 (ES007062), NIEHS R01 (ES028802), and the Michigan Biological Research Initiative on Sex Differences in Cardiovascular Disease (M-BRISC). Data Sharing: All sequencing data will be uploaded to GEO upon acceptance of this manuscript.

## Supplementary Figures

**Figure S1:** UpSet plot showing all significantly enriched GO terms for the DMRs and the number of pathways overlapping across the various groups.

**Figure S2:** Volcano plots showing the number of significantly up or down-regulated genes between weaning (PND21) and 5 months of age, or between 5 months and 10 months of age. Differentially expressed genes were defined as those with a FDR<0.05 and an absolute logFC >2.

**Figure S3:** Venn diagrams showing the number of genes in each sex exhibiting age-related differential methylation and expression, broken out by each of the 3 time point comparisons (weaning vs. 5 months of age, 5 months of age vs. 10 months of age, and weaning vs. 10 months of age).

