## Supplementary figures and images for "Sex-Specific Deflection of Age-Related DNA Methylation and Gene Expression in Mouse Heart by Perinatal Toxicant Exposures"

### Supplemental Figures S1-S3

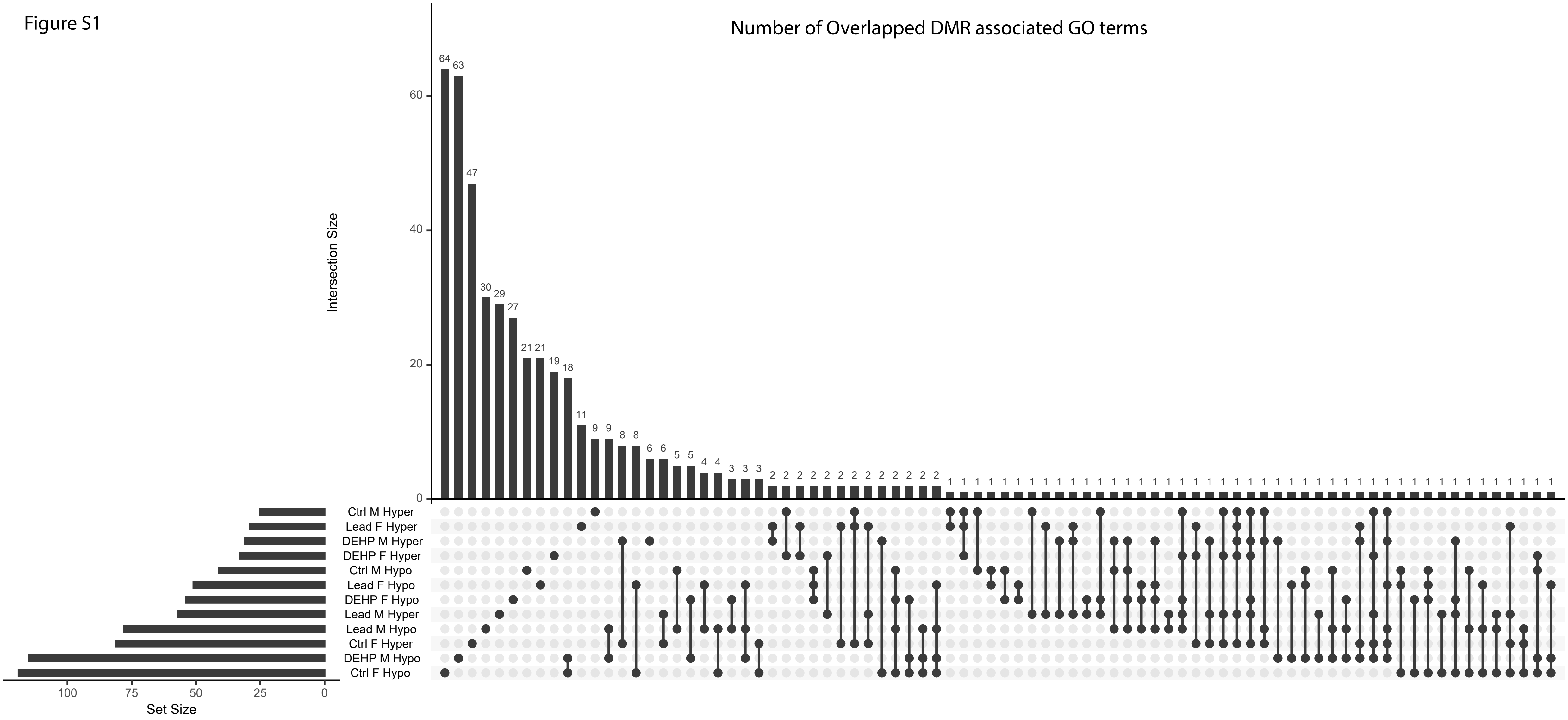

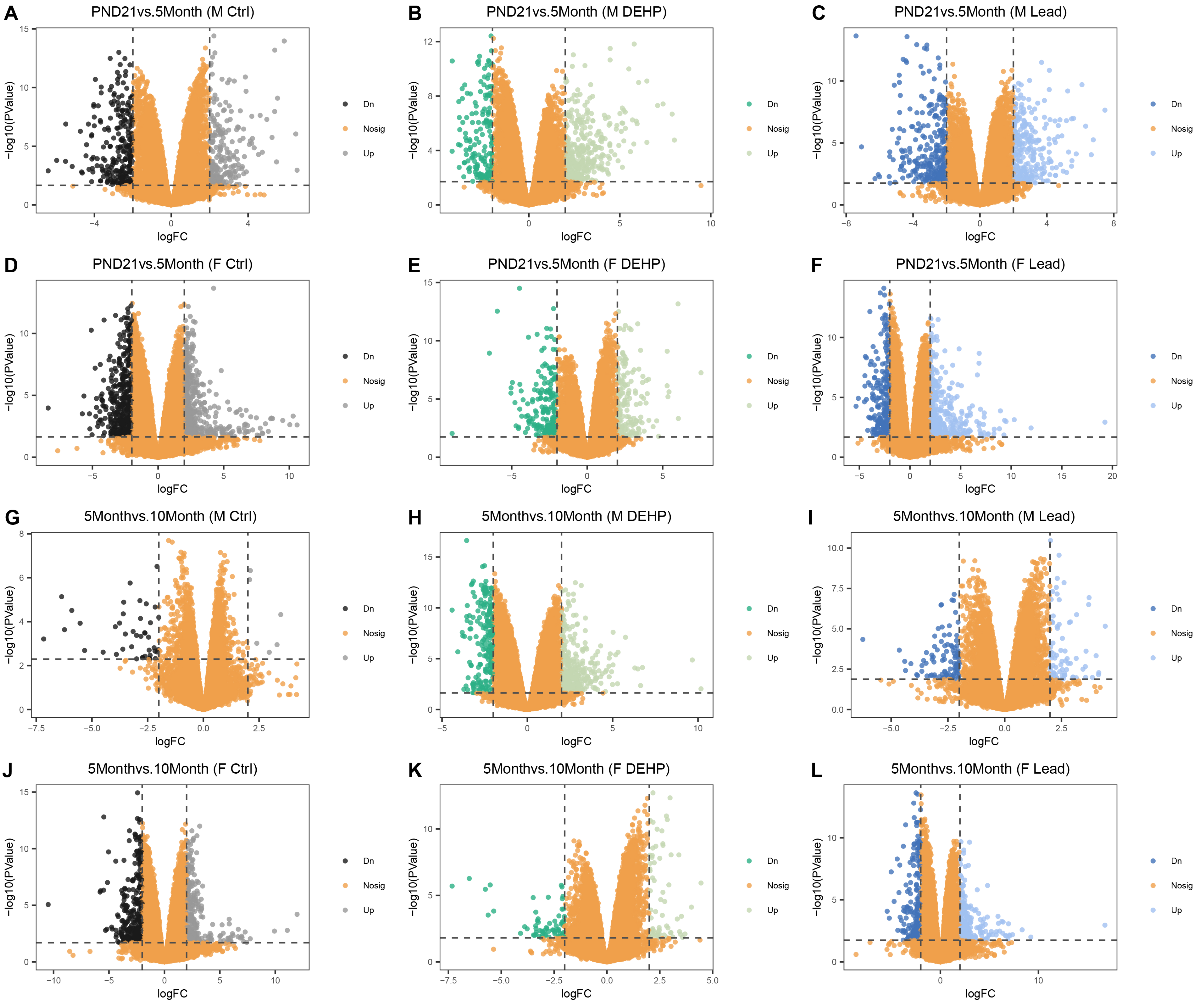

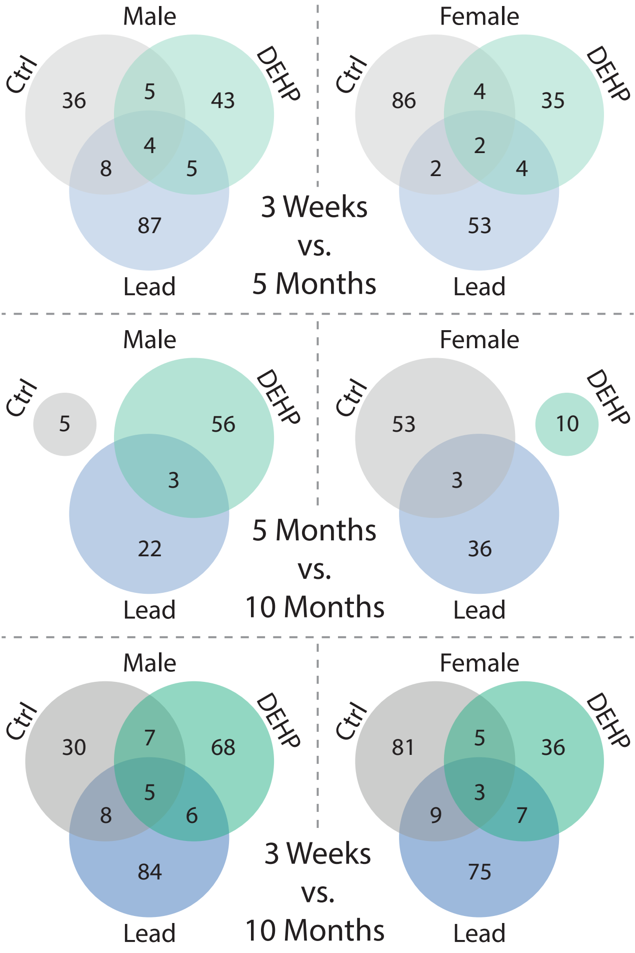
